# HSATII RNA is induced *via* a non-canonical ATM-regulated DNA-damage response pathway and facilitates tumor cell proliferation and movement

**DOI:** 10.1101/2020.05.25.115238

**Authors:** Maciej T. Nogalski, Thomas Shenk

## Abstract

Pericentromeric human satellite II (HSATII) repeats are normally silent, but can be actively transcribed in tumor cells, where increased HSATII copy number is associated with a poor prognosis in colon cancer, and in human cytomegalovirus (HCMV)-infected cells, where the RNA facilitates viral replication. Here, we report that HCMV infection or treatment of ARPE-19 diploid epithelial cells with the DNA-damaging agents, etoposide and zeocin, induced HSATII RNA expression, and a kinase-independent function of ATM was required for the induction. Additionally, various breast cancer cell lines growing in adherent, 2-dimensional cell culture expressed HSATII RNA at different levels, and levels were markedly increased when cells were either infected with HCMV or treated with zeocin. High levels of HSATII RNA expression correlated with enhanced migration of breast cancer cells, and knockdown of HSATII RNA reduced cell migration and the rate of cell proliferation. Our investigation links high expression of HSATII RNA to the DNA damage response, centered on a non-canonical function of ATM, and demonstrates a role for the satellite RNA in tumor cell proliferation and movement.

**SIGNIFICANCE:** HSATII RNA is associated with cancer progression, immunostimulation and, as we recently reported, it plays an important role in herpesvirus infections. However, the understanding of cellular processes responsible for the expression of HSATII RNA has been limited. Our current investigation identified a non-canonical, ATM kinase-independent DNA-damage response pathway as a common cellular mechanism regulating HSATII RNA induction in virus-infected cells or cells treated with DNA-damaging agents. Additionally, our study provides a link between expression of HSATII RNA and the cellular growth and migration phenotypes of cancer cells, establishing a new paradigm to study the biological consequences of HSATII RNA expression, i.e., treatment of normal diploid and tumor cells with DNA-damaging agents.

## INTRODUCTION

Repetitive DNA sequences comprise more than 50% of the human genome (1) and include short (SINE) and long (LINE) interspersed nuclear elements, DNA transposons, long terminal repeat (LTR) transposons and satellite repeats. Even though repetitive DNA elements are ubiquitous in the human genome, there is a relatively limited understanding of their functions and the molecular mechanisms regulating their expression. Satellite DNAs (satDNAs), which account for ∼3% of the genome (1), are constituents of centromeric and pericentromeric heterochromatin, and have been implicated in chromosome organization and segregation, kinetochore formation as well as heterochromatin regulation (2). Next-generation sequencing showed these genomic sites, originally thought to be largely transcriptionally inert, can produce non-coding RNAs (ncRNAs), contributing to satDNA functions (3-6).

Altered patterns of transcription, including the induction of ncRNA accumulation, occur in tumors (7-9). Several ncRNAs originating from satDNA regions of the genome are expressed in cancer cells, such as human alpha-satellite repeat (Alpha/ALR) RNA, human satellite II (HSATII) RNA and its mouse counterpart GSAT RNA (10-13). While some satDNA transcription is stress-dependent (14) or triggered during apoptotic, differentiation or senescence programs in cells (15, 16), HSATII RNA accumulation was found to be refractory to these generalized stressors. Rather, it was induced in a series of colorectal cancer cells when they were grown under non-adherent conditions or as xenografts in mice (17). High expression of Alpha/ALR and HSATII RNAs can lead to their reverse transcription and stable reintegration into the human genome, expanding their genomic copy numbers (17). Elevated copies of genomic HSATII sequences were found in primary human colon tumors and correlated with lower survival rates of colon cancer patients (17). Additionally, HSATII and GSAT RNAs have a nucleotide motif usage that is distinct from that commonly seen in most non-coding transcripts, resulting in their immunostimulatory potential (18).

Viruses de-regulate many biological processes, often in a similar manner to aberrations seen in cancer cells. Those biological changes caused by infections not only resemble cancer, but in some cases contribute to oncogenesis (19). Indeed, it is estimated that 12-20% of all cancers have a viral etiology and often are linked to persistent or chronic viral infections (20-22).

Human cytomegalovirus (HCMV) is a β-herpesvirus that infects a large percentage of the adult population worldwide. Infection in immunocompetent people is typically asymptomatic. In contrast, HCMV is a life-threatening opportunistic pathogen in immunosuppressed individuals (23-25), and a major infectious cause of birth defects (26). Many of the biological changes seen in HCMV-infected cells resemble those common to cancers, including activation of pro-oncogenic pathways, changes in cellular metabolism and increased cell survival (27-32). Given similarities in the molecular symptoms of HCMV infections and tumorigenesis, as well as its reliable presence in glioblastoma tumors, HCMV has been suggested to play a role, perhaps an oncomodulatory role, in the etiology of several human cancers (29, 33-39). However, its high prevalence has made it difficult to establish causality in the disease (40).

Recently, we determined that HCMV infection induces HSATII RNA in fibroblasts, among other satDNA transcripts, to a similar extent as reported for tumor cells (41). Elevated HSATII RNA was also detected in biopsies of CMV colitis, showing that the RNA is elevated in human infections and raising the possibility that elevated HSATII RNA may influence the pathogenesis of HCMV disease. When the amount of HSATII RNA in HCMV-infected cells was reduced by knockdown (KD) with locked nucleic acid (LNA) oligonucleotides, viral gene expression, DNA accumulation and yield were reduced. Moreover, KD experiments revealed that HSATII RNA affects several infected-cell processes, including protein stability, posttranslational modifications and motility of infected cells.

The HCMV IE1 and IE2 proteins induce HSATII RNA in infected cells (41); and are known to regulate the Rb and E2F protein families, which contribute to efficient viral replication in part through induction of a DNA damage response (DDR) (42, 43). Here, we have tested the possibility that these two activities of the viral IE1 and IE2 proteins - induction of a DDR and HSATII RNA - are related. We report that the DNA-damaging agents, zeocin and etoposide, which induce DNA double-strand breaks (DSBs) and a DDR (44, 45), markedly induce HSATII RNA in diploid epithelial cells growing in adherent, 2-dimensional culture. Consistent with this observation, ATM, a key regulator of DDRs, is required for induction of the satellite RNA by infection or DNA-damaging agents. HSATII RNA induction requires a non-canonical, kinase-independent ATM function. Finally, we demonstrate that HSATII RNA is expressed in a variety of breast cancer cell lines grown in adherent culture, the RNA is further induced in these cell lines by HCMV infection or zeocin treatment, and HSATII RNA levels correlate with their migration and proliferation rates. In sum, our data argues that the DDR leads to elevated HSATII RNA levels, potentially enhancing the fitness of both HCMV and tumor cells.

## RESULTS

### The DDR regulates HSATII RNA expression

Ectopic expression of the HCMV IE1 and IE2 proteins can induce HSATII RNA expression in the absence of infection (41). As IE1 and IE2 induce a DDR that supports efficient viral replication (42, 43), we speculated that the DDR could play a role in the expression of human satDNAs.

To test this hypothesis, ARPE-19 cells were stressed by three regimens that induce a DDR: UV-C irradiation, H_2_O_2_ treatment or serum withdrawal (46-48). Exposure of these diploid epithelial cells, which normally express HSATII RNAs at very low basal levels, to these stressing agents induced HSATII RNA to a very limited extent, 2-3 fold (Fig. 1*A*), in contrast to the 100-1000-fold induction observed in response to virus infection (41). As a proxy to monitor successful DDR activation, the induction of p53 activity was monitored by assaying the levels of p53-dependent cyclin-dependent kinase inhibitor 1 (p21; encoded by the CDKN1A gene) (46) and p53-regulated Damage Induced Noncoding (DINO) lncRNA (49). As expected, both were elevated in cells exposed to all three treatments (Fig. 1*A*).

**Fig. 1.**
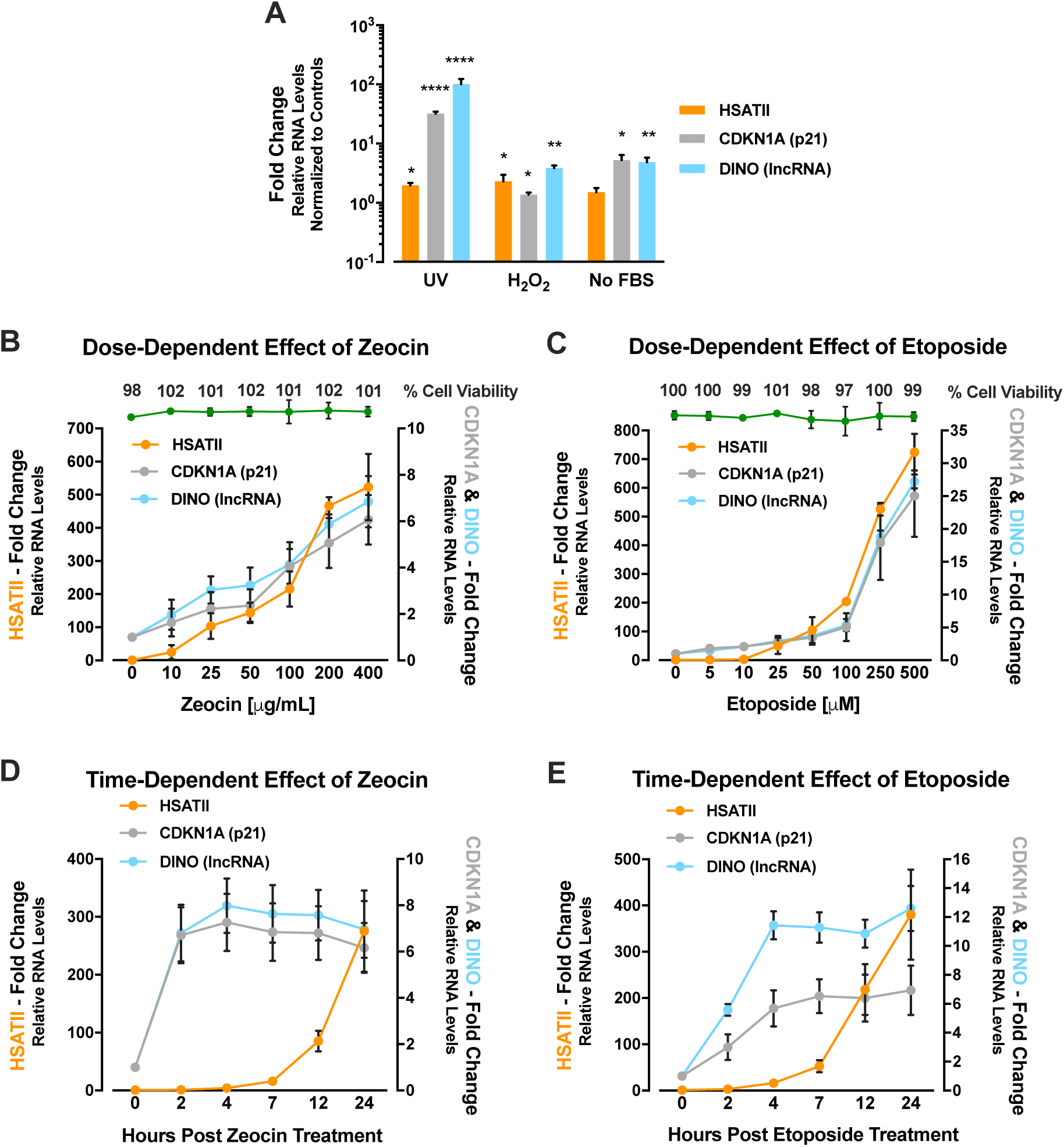
DNA damage response regulates HSATII RNA expression. (*A*) Limited induction of HSATII RNA by UV, H_2_O_2_, or serum withdrawal. ARPE-19 cells were exposed to UV-irradiation (50 J/m^2^), H_2_O_2_ (100 nM) or serum withdrawal (no FBS). RNA samples were collected at 24 hpt. RT-qPCR was performed to quantify HSATII, CDKN1A or DINO transcripts, using GAPDH as an internal control. Data are presented as a fold change (mean ± SD; n=3). \**P*<0.05, \*\**P*<0.01, \*\*\**P*<0.001. (*B* and *C*) Induction of HSATII RNA by treatment with DNA-damaging drugs. ARPE-19 cells were treated with increasing concentrations of zeocin (*B*) or etoposide (*C*) with H_2_O or DMSO, respectively, as solvent controls. RNA samples were collected at 24 hpt, and RT-qPCR was performed to quantify HSATII, CDKN1A and DINO transcripts, using GAPDH as an internal control. Data are presented as a fold change (mean ± SD; n=3). Insert: At 24 hpt, cell viability was assessed at each indicated drug concentrations. Data is presented as % viable cells, (mean ± SD; n = 3). (*D* and *E*) Drug-induced HSATII RNA accumulation occurs after a delay. ARPE-19 cells were treated with zeocin (200 μg/mL) (*D*), etoposide (200 μM) (*E*) and H_2_O or DMSO, respectively, as solvent controls. RNA samples were collected at indicated times. RT-qPCR was performed to quantify HSATII, CDKN1A and DINO transcripts, using GAPDH as an internal control. Data are presented as a fold change (mean ± SD; n=3).

We reasoned that the failure of these treatments to induce HSATII RNA might be a consequence of the specific type of DNA damage and the resulting response that they induce. For example, while UV-C irradiation can lead to DNA double-strand brakes (DSBs), the specific role of UV-C in DNA damage depends on the replicative state of treated cells, as those breaks usually arise from the replication of unrepaired UV-induced DNA lesions (50, 51). It was possible that UV-C and the other treatments were not efficient in inducing HSATII RNA. Therefore, we tested the effects of zeocin and etoposide. Etoposide, similarly to doxorubicin, is an anticancer agent that poisons topoisomerase II, stabilizing DNA-protein complexes, which ultimately leads to stalled replication forks (52). Zeocin belongs to the bleomycin/phleomycin-family of antibiotics and, as a radiomimetic, has similar effects on DNA as ionizing radiation (53). Both drugs induce DNA DSBs (44, 45), and etoposide has been shown to stimulate the expression and retrotransposition of another repeated element, mouse B2 SINE RNA (54, 55). ARPE-19 cells were exposed to increasing concentrations of zeocin or etoposide and HSATII, CDKN1A and DINO RNA expression were monitored. Zeocin caused a rapid, dose-dependent induction of HSATII RNA, reaching an ∼500-fold increase relative to untreated cells, at the maximum dose tested (400 μg/mL) (Fig. 1*B*). Etoposide also elevated HSATII RNA levels, reaching an ∼700-fold increase at the highest dose tested (500 μM) (Fig. 1*C*). CDKN1A and DINO RNA expression mirrored HSATII RNA accumulation, reaching a 6-7-fold induction in cells exposed to 400 μg/mL of zeocin (Fig. 1*B*) and ∼25-fold induction in cells treated with 500 μM of etoposide (Fig. 1*C*). No ARPE-19 cell toxicity was detected at any drug concentration after a 24 h treatment (Figs. 1*B* and *C*).

The kinetics of HSATII, CDKN1A and DINO RNA expression were monitored by RT-qPCR from 0-24 h post treatment (hpt) with zeocin (200 μg/mL) or etoposide (200 μM). HSATII RNA expression started to increase between 7-12 hpt with either zeocin or etoposide, reaching a 300-fold induction at 24 hpt compared to solvent control-treated cells (Figs. 1*D* and *E*). The dynamics of CDKN1A and DINO RNA expression were accelerated compared to HSATII RNA expression, as both transcripts reached their maximum levels within the first 2-4 hpt, suggesting a possible decoupling of DDR/p53-dependent pathways from the mechanism regulating HSATII RNA induction.

### Zeocin treatment substantially mimics the transcriptional effect of HCMV infection on HSATII RNA expression

To compare the induction HSATII RNA by virus versus DNA-damaging agent, ARPE-19 cells were infected with HCMV or treated with zeocin. The infection used an epithelial cell-grown HCMV strain (TB40/E-GFP-epi) that efficiently infects ARPE-19 cells (56). Mock-infection and solvent treatment served as controls. After 24 h, RNA was isolated and analyzed by using RNA-seq. By analyzing only uniquely mapped HSATII reads, RNA-seq analysis confirmed that zeocin effectively induces HSATII expression in ARPE-19 cells (Fig. 2*A*). Moreover, as in HCMV-infected cells (41), HSATII RNA in zeocin-treated cells is produced preferentially from chromosome 1, 2, 10, and 16. We detected a higher prevalence of HSATII-specific reads from chromosome 1 in cells exposed to zeocin (∼15%) compared to only ∼5% of these reads in samples from HCMV-infected cells (Fig. 2B). However, in both HCMV- or zeocin-treated cells the majority (∼86% or 76%, respectively) of HSATII specific reads mapped to chromosome 16 (Fig. 2*B*).

**Fig. 2.**
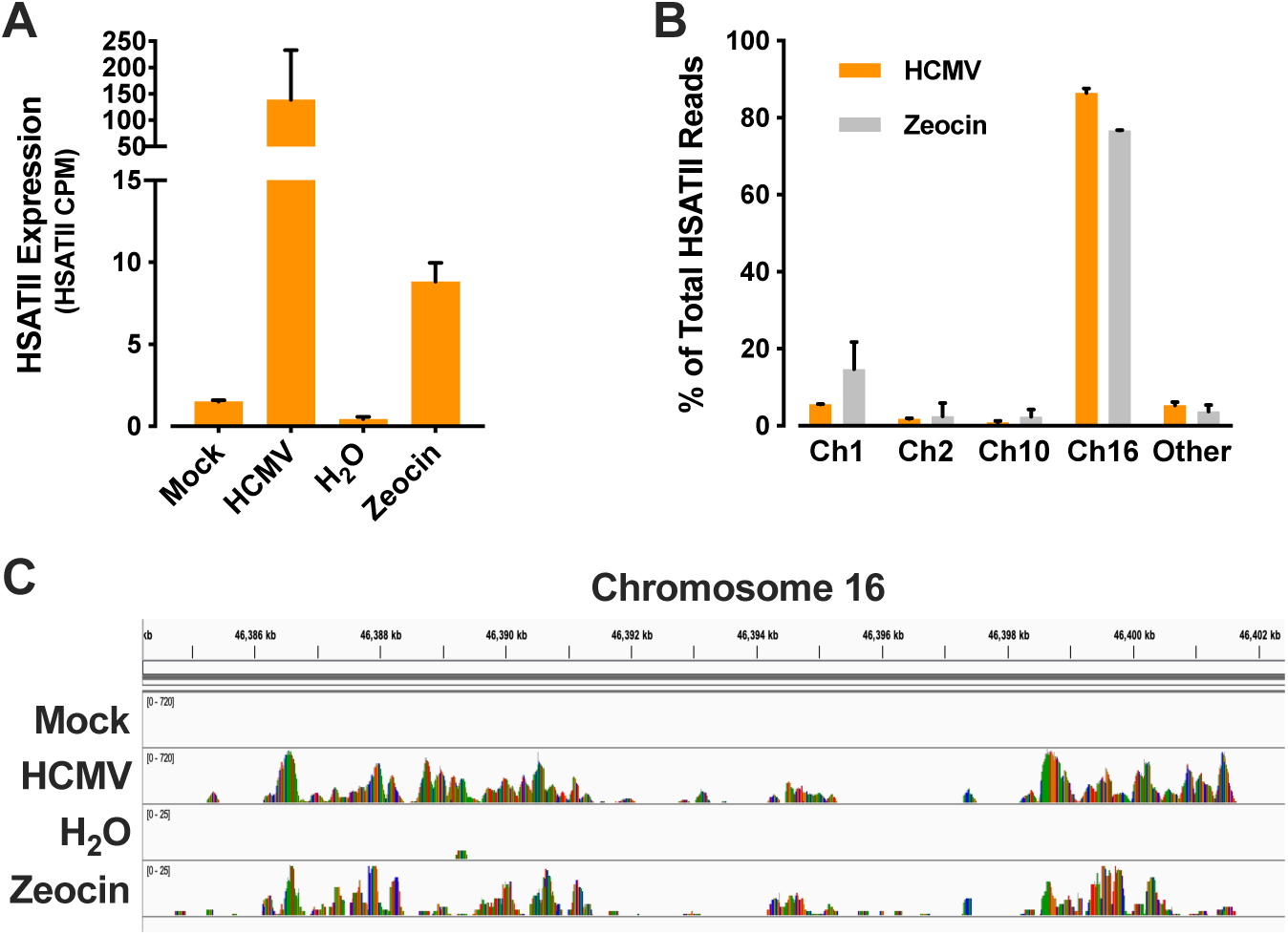
Zeocin treatment partially mimics the transcriptional effect of HCMV infection on HSATII RNA expression. ARPE-19 cells were infected with HCMV (1 TCID50/cell), or treated with zeocin (100 μg/mL) or H_2_O as a solvent control, and RNA samples were collected at either 24 hpi or 24 hpt, respectively. RNA was isolated and analyzed using RNA-seq. (*A*) Quantification of HSATII RNA induction by RNA-seq analysis. HSATII RNA expression in terms of counts per million reads mapped (CPM) was computed and normalized across samples (n=2). (*B*) Chromosomal distribution of HSATII RNA expression. HSATII chromosomal origin in HCMV-infected or zeocin-treated cells was depicted based on the number of unique HSATII reads mapped to specific chromosomal loci. Data are presented as a percentage of total HSATII reads (mean ± SD. n=2). (*C*) Example loci of HSATII RNA expression on chromosome 16. HSATII read coverage for HCMV-infection versus zeocin treatment was based on reads mapped to human genome and read coverage track at the HSATII locus on chromosome 16 was visualized using IGV.

Read coverage tracks at the HSATII-rich locus on chromosome 16 were mapped with the Integrative Genomics Viewer (IGV) using STAR aligner (57)-created BAM files (Fig. 2*C*). The data from mock-infected and solvent-treated cells showed very few, if any, reads mapped to this HSATII locus (Fig. 2*C*). The read coverage tracks based on RNA-seq data from HCMV-infected or zeocin-treated ARPE-19 cells display a similar distribution in this HSATII–rich locus spanning approximately 17 kb of DNA. Therefore, these data strengthened our view that common mechanisms might be responsible for the induction of HSATII RNA in both HCMV-infected and DNA-damaging drug-treated cells.

### Zeocin treatment partially mimics the HCMV-induced transcriptome

To assess common biological processes regulated by infection and drug, ARPE-19 cells were HCMV-infected or zeocin-treated, and, again, mock-infection or solvent treatment served as controls. The RNA-seq analysis determined that 654 and 914 cellular RNAs were significantly (q < 0.05) modulated in HCMV-infected and zeocin-treated cells, respectively, and 87 RNAs were modulated by both treatments (Fig. 3*A*). The 87 co-regulated genes were further analyzed by Gene Set Enrichment Analysis (GSEA) (58) using the hallmark gene set of the Molecular Signatures Database (MSigDB) (59) and found to be strongly associated with multiple gene subsets related to DNA damage/repair and other processes, which, if deregulated, associate with enhanced cancer progression (60-63) (Fig. 3*B*). As we were interested in determining the common molecular mechanisms that govern HSATII RNA induction in infected and zeocin-treated cells, we focused on the role of HCMV- and drug-induced DDR pathways in HSATII RNA expression and its possible effect on cancer cell biology.

**Fig. 3.**
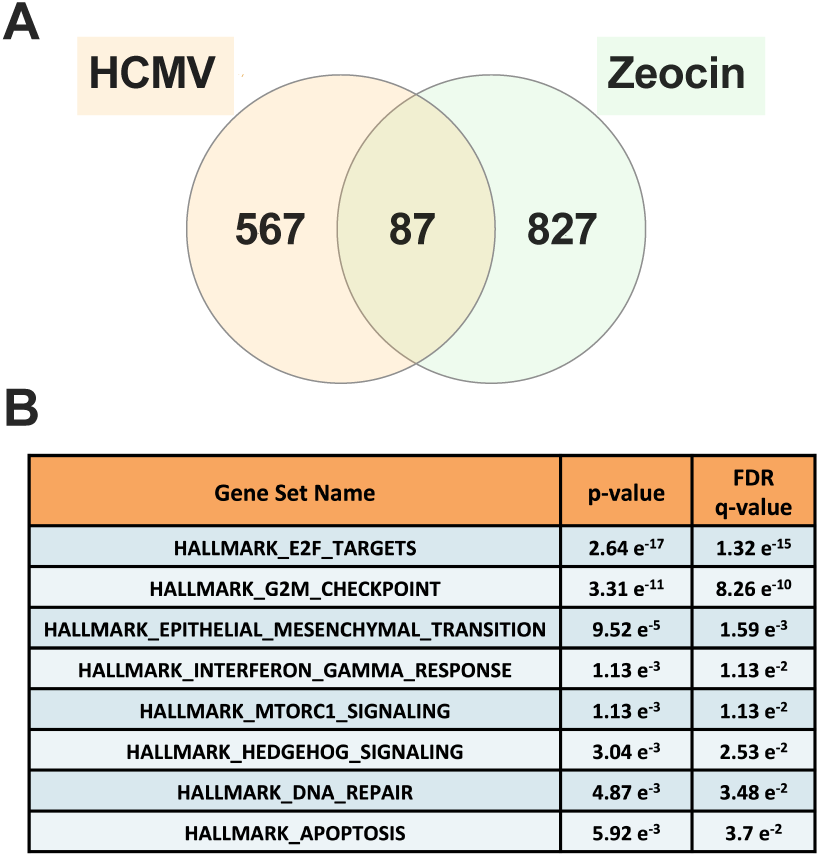
Zeocin treatment partially mimics the HCMV-induced transcriptome. ARPE-19 cells were mock- or TB40/E-GFP-epi-infected (1 TCID_50_/cell), or treated with zeocin (100 μg/mL) or H_2_O as a solvent control. After 24 h, RNA samples were collected, RNA was isolated and analyzed using RNA-seq. (*A*) Cellular genes modulated by both HCMV and zeocin. Venn diagram showing significantly expressed genes (q<0.05) either in HCMV-infected or zeocin-treated cells vs. respective controls. (*B*) Categories of gene sets modulated by both HCMV and zeocin. GSEA was performed on the genes differentially expressed in both HCMV-infected and zeocin-treated cells using the hallmark gene set of MSigDB. Identified specific gene set names are categorized based on increasing *p*-value and FDR q-value.

### ATM is required for HSATII RNA expression

Different types of DNA damage govern the activation of DDR pathways through key regulators: ATM, ATR and DNA-PKcs (64-66), and each of these PI3-kinases, when activated, phosphorylates the histone H2A family member, H2AX, generating γ-H2AX (67). HCMV infection induces accumulation of γ-H2AX (42, 43) as does zeocin (68). To confirm that both agents induce this marker in ARPE-19 epithelial cells, untreated, HCMV-infected, or zeocin-treated (100 μg/mL) cells were fixed after 24 h, stained with antibody recognizing γ-H2AX protein and representative images were captured. Both HCMV-infected and zeocin-treated cells had elevated γ-H2AX signals (Fig. 4*A*), however, cells exposed to zeocin were characterized by a higher number of γ-H2AX-positive cells (65%) than HCMV-infected cells (14%) at 24 h after infection or drug treatment (Fig. 4*B*).

**Fig. 4.**
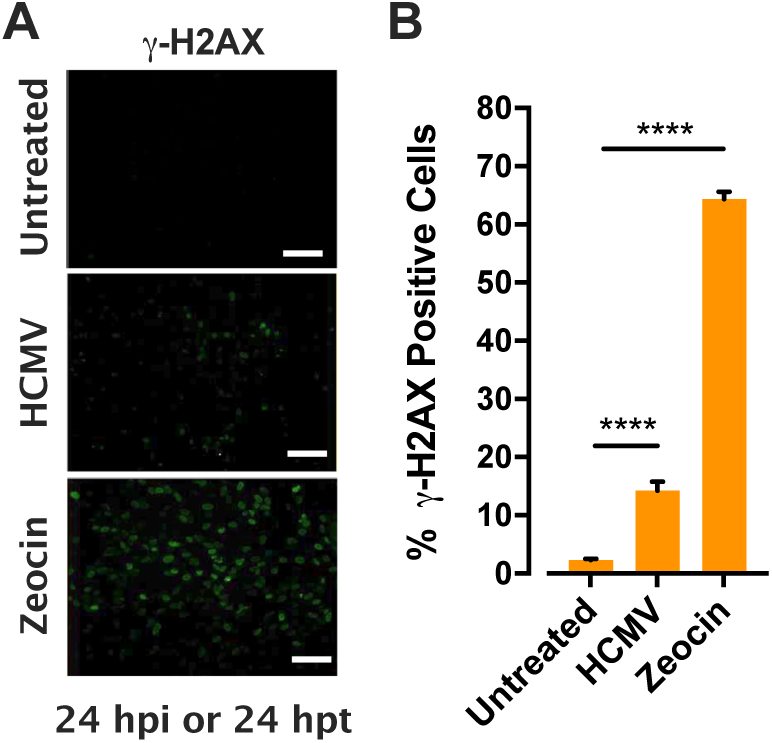
HCMV and zeocin induce γ-H2AX in ARPE-19 cells. Cells were mock- or TB40/E-GFP-epi-infected (3 TCID50 /cell) or treated with zeocin (200 μg/mL). After 24 h, cells were stained for γ-H2AX, nuclei were counterstained with the Hoechst stain, cells were visualized (*A*: scale bars=100 µm) and % γ-H2AX-positive cells were quantified (*B*: mean ± SD; n=3). \*\*\*\**P*<0.0001.

We next scrutinized possible contributions of ATM, ATR and/or DNA-PKcs to HSATII RNA accumulation. ARPE-19 cells were transfected with NT siRNA or siRNAs targeting ATM, ATR or PRKDC (encodes DNA-PKcs) transcripts. 48 h later, the cells were infected with HCMV or treated with zeocin (100 μg/mL) for 24 h. All siRNAs efficiently reduced their targeted transcripts (75-90% decrease) (Fig. 5*A*). However, only ATM-specific siRNA treatment led to a significant reduction of HSATII RNA levels (∼70% decrease) in HCMV-infected (Fig. 5*A*, left panel) or zeocin-treated cells (Fig. 5A, right panel). To directly assess the importance of ATM kinase activity (66, 69, 70) in HSATII RNA induction the effect of Ku-55933 or AZ31, ATM kinase inhibitors (71, 72), were tested. Neither drug compromised the viability of ARPE-19 cells when tested at concentrations up to 50 μM (Fig. 5B, left panel). However, because we noticed morphological changes in treated cells at the highest dose (50 μM), we used Ku-55933 and AZ31 at 20 μM, which is in the range of concentrations at which these drugs are typically used (71-73). Cells were pretreated with Ku-55933 or AZ31 for 2 h before being treated with zeocin (100 μg/mL). Interestingly, at 24 hpt neither ATM kinase inhibitor affected HSATII RNA levels (Fig. 5*B*, right panel). However, both kinase inhibitors negatively (∼70% decrease) affected expression of CDKN1A and DINO transcripts when compared to DMSO-treated control cells, demonstrating that the drugs were active.

**Fig. 5.**
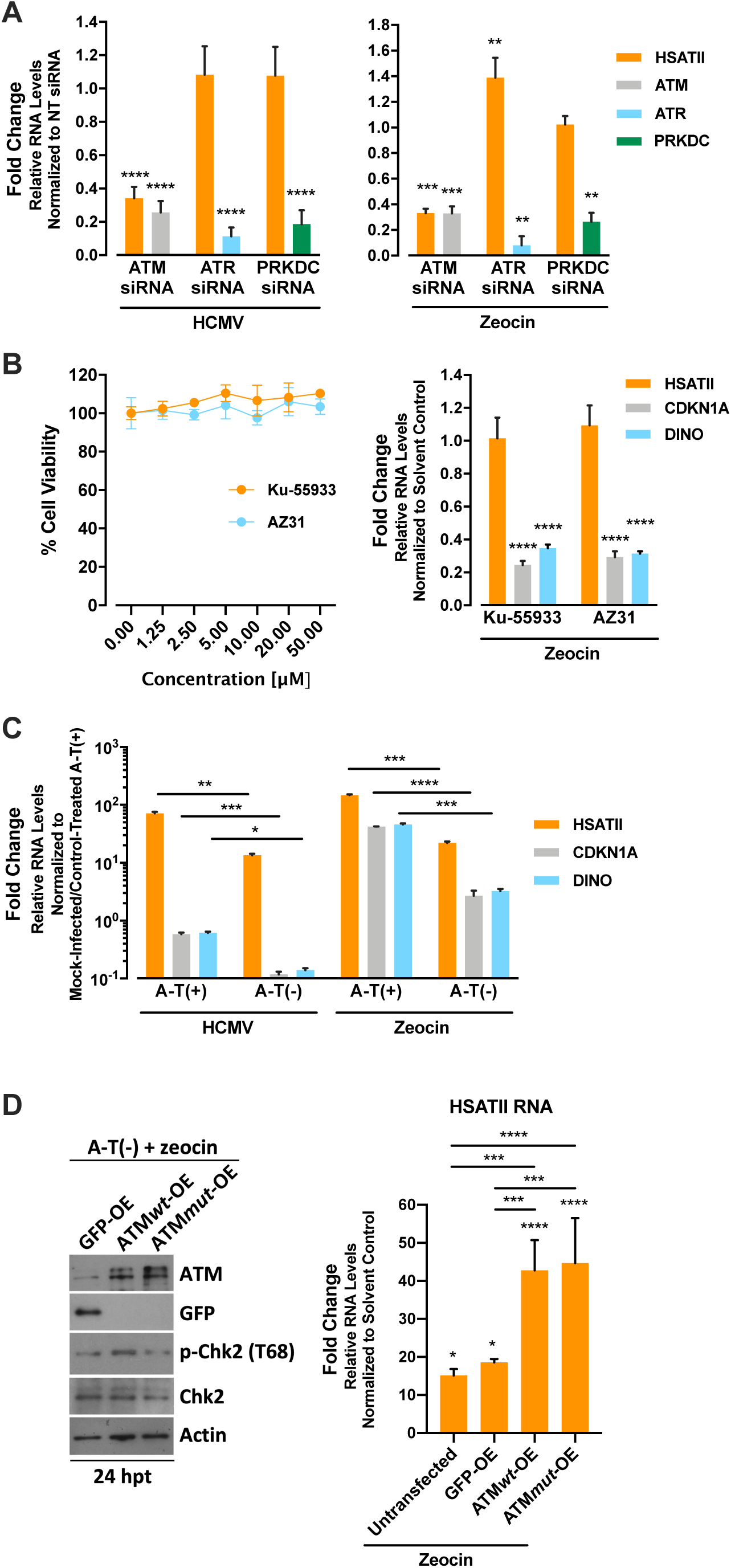
ATM-based DNA damage response regulates HSATII RNA expression. (*A*) ATM KD inhibits the induction of HSATII RNA by HCMV infection or zeocin treatment. ARPE-19 cells were transfected with ATM-, ATR- or PRKDC-specific siRNAs or NT siRNA as a control. After 48 h, cells were infected with TB40/E-GFP-epi (1 TCID_50_/cell) or zeocin (100 μg/mL). RNA samples were collected 24 h later. RT-qPCR was performed to quantify HSATII, ATM, ATR and PRKDC transcripts, using GAPDH as an internal control. Data are presented as a fold change (mean ± SD; n=3-10). (*B*) HSATII RNA is induced less in A-T(-) cells than in A-T(+). A-T(+) and A-T(-) cells were uninfected, infected with TB40/E-GFP (1 TCID_50_/cell), or treated with zeocin (100 μg/mL) or H_2_O as a solvent control. RNA samples were collected 24 h later. RT-qPCR was performed to quantify HSATII, CDKN1A and DINO transcripts, using GAPDH as an internal control. Data are presented as a fold change (mean ± SD; n=3). (*C*) ATM kinase inhibitors do not block the induction of HSATII RNA by zeocin. Left panel: ARPE-19 cells were treated with various concentrations of Ku-55933 or AZ31. At 24 hpt, cell viability was assessed. Data is presented as % viable cells, (mean ± SD; n=3). Right panel: ARPE-19 cells were treated with DMSO, Ku-55933 (20 μM) or AZ31 (20 μM) for 2 h before zeocin (100 μg/mL) was added. RNA samples were collected 24 hpt, and RT-qPCR was performed to quantify HSATII, CDKN1A and DINO transcripts, using GAPDH was used as an internal control. Data are presented as a fold change (mean ± SD; n=3). (*D*) Ectopic expression of kinase-dead ATM (ATM*mut*) can enhance the induction of HSATII RNA in A-T(-) cells by zeocin. Left panel: A-T(-) cells were transfected with GFP-, ATM*wt*- or ATM*mut*-expressing plasmids. After 48 h, cells were exposed to zeocin (100 μg/mL) and protein samples were collected 24 h later. Proteins were assayed by Western blot analysis performed using antibodies recognizing ATM, GFP, phospho-Chk2 (T68), Chk2 and actin as a loading control. Right panel: A-T(-) cells were transfected with GFP-, ATM*wt*- or ATM*mut*-expressing plasmids and untransfected cultures served as controls. After 48 h, cells were treated with zeocin (100 μg/mL) and RNA samples were collected 24 h later. RT-qPCR was performed to quantify HSATII. GAPDH was used as an internal control. Data are presented as a fold change (mean ± SD. n=4). \**P*<0.05, \*\**P*<0.01, \*\*\**P*<0.001, \*\*\*\**P*<0.0001.

To further establish a role for ATM in HSATII RNA expression, we used ataxia-telangiectasia fibroblasts, A-T(-) cells, that carry a missense mutation in the carboxy-terminal PI3-kinase-like domain of ATM, and contain very low levels of mutant protein (74, 75). A derivative of this cell line, A-T(+) cells, with a stably expressed, functional ATM gene (75) was used as a control. Cells were HCMV-infected or treated with zeocin (100 μg/mL). Mock infection and solvent treatment served as controls. After 24 h, infected and drug-treated A-T(+) cells were characterized by a 71x or 137x induction of HSATII RNA, respectively; and zeocin-treated, but not HCMV-infected, cells contained higher levels of CDKN1A and DINO RNA. HSATII RNA was also induced in A-T(-) cells following HCMV infection or zeocin treatment, but to an ∼6-fold lower level when compared A-T(+) cells (Fig. 5*C*). Both treatments resulted in significantly lower expression of CDKN1A and DINO RNAs (∼15x and ∼5x lower, respectively) in A-T(-) cells when compared to A-T(+) cells. Together these data further implicate ATM in HSATII RNA expression following infection or treatment with DNA-damaging drugs.

Next, to confirm a kinase-independent role for ATM in HSATII RNA expression, we ectopically expressed wild-type ATM (ATM*wt*), kinase-dead ATM (ATM*mut*) or GFP, as a control, in the A-T(-) fibroblasts and followed with zeocin treatment. As originally demonstrated by Ziv *et al*. A-T(-) fibroblasts are characterized by a very low level of ATM protein (75) and transfection of the cells with ATM-encoding plasmids resulted in the efficient expression of ATM*wt* or ATM*mut* (Fig. 5D, left panel). ATM*wt-*but not ATM*mut*-expressing cells responded to zeocin treatment with enhanced phosphorylation of Chk2 at threonine 68, as predicted. HSATII RNA levels were monitored in these cells. Similarly to the data presented in figure 5*C*, zeocin treatment increased HSATII expression in untransfected or GFP A-T(-) cells (Fig. 5*D*, right panel). Importantly, however, both ATM*wt*- and ATM*mut*-expressing A-T(-) cells responded to zeocin treatment with similarly increased HSATII RNA expression, when compared to untransfected or GFP-expressing cells. This experiment confirms that ATM regulates HSATII expression, but independently from its kinase activity.

### HSATII RNA expression is induced by ATM *via* a Chk1/2- and p53-independent pathway

ATM and ATR phosphorylate two downstream kinases, checkpoint 1 and 2 (Chk1 and Chk2 encoded by the CHEK1 and CHEK2 genes, respectively), which ultimately regulate p53 activity in a context-dependent manner (66, 69, 70). Chk2 is normally activated by ATM-mediated phosphorylation, so we tested whether it is required for accumulation of HSATII RNA. As a control, we also tested Chk1. 48 h prior to HCMV infection or treatment with zeocin (100 μg/mL), ARPE-19 cells were transfected with NT siRNA or siRNA specifically targeting CHEK1 or CHEK2 transcripts (encoding Chk1 or Chk2, respectively). The treatment resulted in a robust decrease in these transcripts at 24 hpi or 24 hpt (Fig. 6*A*). However, we did not detect a statistically significant decrease of HSATII RNA levels in CHEK1 or CHEK2 KD cells upon HCMV infection or zeocin treatment, consistent with the lack of a requirement for ATM kinase activity (Fig. 5), and further arguing that the canonical ATM-Chk2 signaling cascade does not play an important role in the expression of HSATII RNA.

**Fig. 6.**
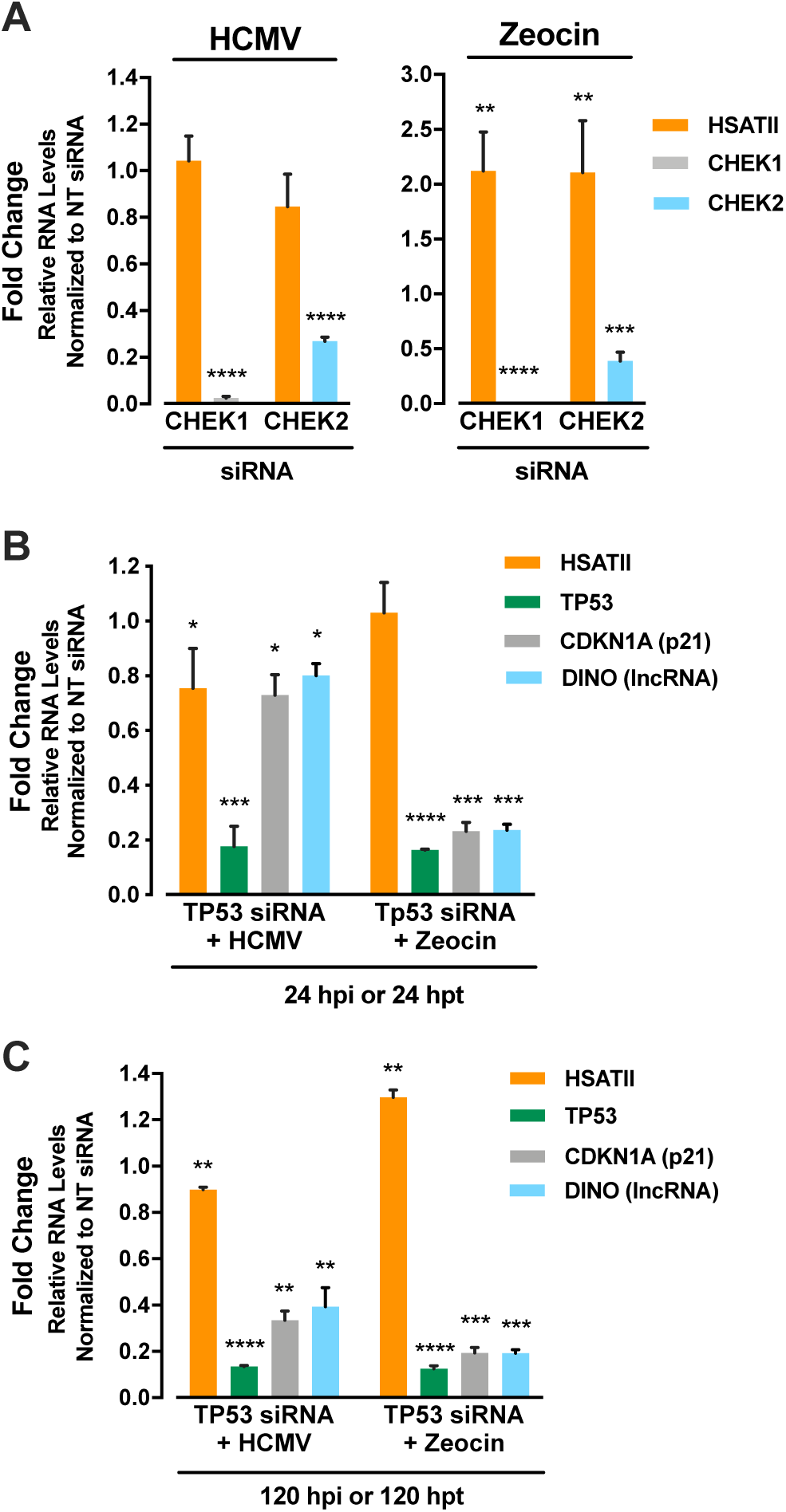
HSATII RNA expression is induced via a Chk1/2- and p53-independent pathway. (*A*) CHEK1 or CHEK2 KD doesn’t block HSATII RNA induction by infection or zeocin. ARPE-19 cells were transfected with CHEK1- or CHEK2-specific siRNA or NT siRNA as a control. After 48 h, cells were infected with TB40/E-GFP-epi (1 TCID_50_/cell) or zeocin (100 μg/mL). RNA samples were collected 24 h later. RT-qPCR was performed to quantify HSATII, CHEK1 and CHEK2 transcripts, using GAPDH as an internal control. Data are presented as a fold change (mean ± SD; n=4). (*B* and *C*) p53 KD does not block HSATII RNA induction following HCMV infection or zeocin treatment. ARPE-19 cells were transfected with TP53 siRNAs or NT siRNA as a control. After 48 h, cells were infected with TB40/E-GFP-epi (1 TCID/cell) or zeocin (100 μg/mL), and RNA samples were collected 24 h (*B*) or 120 h (*C*) later. RT-qPCR was performed using specific primers to HSATII, TP53, CDKN1A or DINO transcripts. GAPDH was used as an internal control. Data are presented as a fold change (mean ± SD; n=3-4. \**P*<0.05, \*\**P*<0.01, \*\*\**P*<0.001, \*\*\*\**P*<0.0001.

Since ATM (76) and Chk2 (77) both phosphorylate and stabilize/activate p53, we also investigated the possible involvement of p53 in HSATII RNA induction. Control and p53 KD ARPE-19 cells were HCMV-infected or treated with zeocin (100 μg/mL), and HSATII, CDKN1A and DINO RNA expression were assessed at 24 (Fig. 6*B*) and 120 hpi/hpt (Fig. 6*C*). TP53-specific siRNA strongly decreased the targeted p53 transcript at both times tested (∼83% and 87%), when compared to NT siRNA-treated cells. p53 KD had a modest effect (10% decrease) on HSATII RNA expression at 120 hpi when compared to control cells, but zeocin treatment either had no effect (24 hpt) or slightly increased HSATII RNA expression (120 hpt). In control experiments, p53 KD had little effect on CDKN1A and DINO expression at 24 h post HCMV infection, but significantly decreased (∼65%) both transcripts at 120 hpi. In comparison, p53 KD strongly (80% decreased) affected CDKN1A and DINO expression at both 24 and 120 h post zeocin treatment.

In sum, ATM protein is required for HSATII RNA accumulation in response to infection or zeocin treatment. However, ATM kinase activity and two downstream elements of the DDR response that are normally activated by ATM-mediated phosphorylation, Chk2 and p53, are not required for HSATII RNA expression.

### Breast cancer cells differ in HSATII RNA levels and sensitivity to HCMV infection and zeocin treatments

HSATII RNA expression has been detected in several cancers, including breast (10, 12), and HSATII copy number gains are a common and negative prognostic feature of colorectal tumors (17). Given the induction of HSATII RNA by HCMV infection (41) and DNA-damaging agents (Fig. 1), and knowing that deregulated DDRs allow cancer cells to achieve high proliferation rates while making them more susceptible to DNA-damaging agents (53, 78), we explored the consequences of induced HSATII RNA expression in breast cancer cells. Initially, we assessed HSATII RNA expression levels in breast cancer cell lines that differed by p53 status, hormonal dependency and invasiveness (Fig. 7*A*). We tested less invasive tumor cell lines: MDA-MB-175VII (ER+, PR-, HER2-, p53 wt), MDA-MB-361 (ER+, PR+/-, HER2-, p53 wt), MCF7 (ER+, PR+, HER2-, p53 wt); more invasive triple negative breast cancer (TNBC) cell lines with mutated p53: SUM1315M02, MDA-MB-231, BT-549; and “normal” MCF-10A breast epithelial cells as well as “normal” diploid ARPE-19 cells (79). Cells were cultured in an adherent, 2-dimensional format using identical media conditions across all lines to prevent variations in results potentially caused by different culture conditions. Upon reaching 80-90% confluency, RNA samples were collected. The basal levels of HSATII RNA in the three less invasive cell lines (MCF-7, MDA-MB-175VII and MDA-MB-361) were 5-6-fold higher when compared to HSATII expression in ARPE-19 cells; and the three more invasive cell lines (SUM1315M02, MDA-MB-231, BT-549) were characterized with 14- to 53-fold higher levels of HSATII RNA compared to less invasive breast cancer cells or ARPE-19 cells. The “normal” MCF-10A breast epithelial cells had HSATII RNA levels similar to the less invasive group of tumor cells, suggesting that the basal level of HSATII RNA might be influenced by the tissue origin of tested cells.

**Fig. 7.**
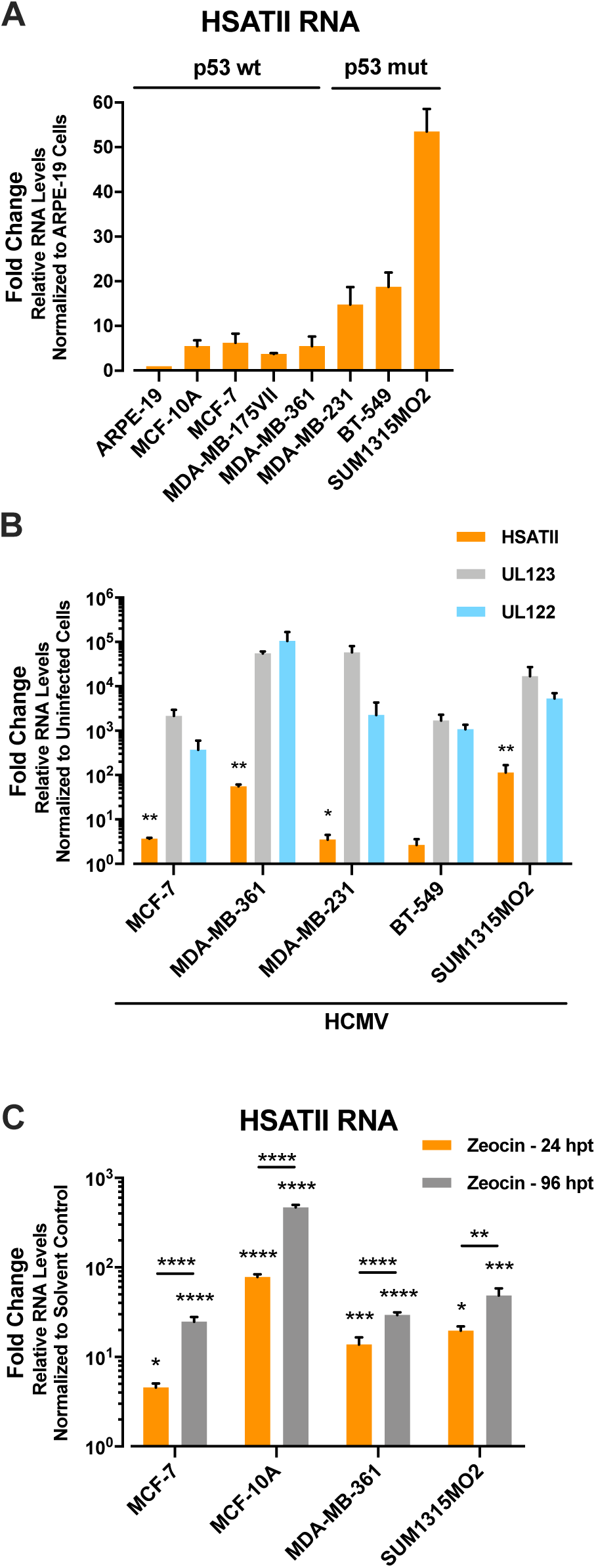
Breast cancer cells differ in HSATII RNA levels and sensitivity to HCMV infection and zeocin treatment. (*A*) Breast cancer cell lines contain HSATII RNA when grown in adherent (2-dimensional) culture. Cell lines were grown to ∼80% confluency and RNA samples were collected. RT-qPCR was performed to quantify HSATII RNA, using GAPDH as an internal control. Data are presented as a fold change (mean ± SD; n=3). (*B*) HCMV infection elevates HSATII RNA levels in breast cancer cell lines. MCF-7, MDA-MB-361, MDA-MB-231, BT-549 and SUM1315MO2 cells were infected with TB40/E-GFP-epi (4 TCID_50_/cell) and RNA samples were collected at 24 hpi. RT-qPCR was performed to quantify HSATII RNA, using GAPDH as an internal control. Data are presented as a fold change (mean ± SD; n=3-5). (*C*) Zeocin treatment elevates HSATII RNA levels in breast cancer cell lines. MCF-7, MCF-10A, MDA-MB-361 and SUM1315MO2 cells were treated with zeocin (200 μg/mL). RNA samples were collected at 24 and 96 hpt and RT-qPCR was performed to quantify HSATII RNA, using GAPDH as an internal control. Data are presented as a fold change (mean ± SD; n=3). \**P*<0.05, \*\**P*<0.01, \*\*\**P*<0.001, \*\*\*\**P*<0.0001.

Next, we tested the induction of HSATII RNA in a selection of the breast cancer cell lines in response to HCMV infection (Fig. 7*B*). At 48 hpi, all tested cells, except BT-549 cells, demonstrated significantly elevated but varied levels of HSATII RNA (from ∼5-fold increase in MCF-7 and MDA-MB-231 to ∼100-fold increase in MDA-MB-361 and SUM1315MO_2_) compared to its levels in uninfected cells. As we previously established that HCMV IE1 and IE2 proteins collaborate to induce HSATII RNA (41) and knowing that HCMV replication is limited or abortive in cancer cells (56, 80, 81), we monitored expression of UL123 and UL122 viral transcripts, encoding the IE1 and IE2 proteins. UL123 and UL122 RNA expression was detected in the breast cancer cell lines upon HCMV infection (Fig. 7B). However, our data did not detect any correlation between levels of HSATII RNA and viral transcripts.

We also tested the effect of zeocin treatment on HSATII RNA induction in these breast cancer cell lines (Fig. 7*C*). At 24 and 96 hpt, all cell lines responded to zeocin with significantly elevated levels of HSATII RNA compared to its levels in solvent control-treated cells. Additionally, a longer exposure (96 h) of cells to the DNA-damaging drug correlated with even higher expression of HSATII RNA. At 96 hpt HSATII RNA levels increased by as much as 240-fold in the case of MCF-10A cells.

### DNA damage-induced HSATII RNA enhances motility and proliferation of breast cancer cells

We previously reported that HSATII RNA KD reduced the motility of HCMV-infected ARPE-19 cells (41). Since HCMV infection and zeocin treatment elevated HSATII RNA expression in breast cancer cell lines (Fig. 7*B* and *C*), we tested whether induced HSATII RNA influences migration of these cells. Infection of breast cancer cells at a high multiplicity with HCMV leads to viral protein expression in only a portion of the cells (56), and this would confound interpretation of the migration assay. Therefore, we treated ARPE-19, MCF-7 and SUM1315MO2 cells with zeocin (200 μg/mL) or H_2_O as a solvent control to guarantee a uniform effect across the cell population (Fig. 8*A*). At 24 hpt, transwell migration assays (82) were performed using equal numbers of cells and cell motility was assessed after an additional 24 h. ARPE-19 and MCF-7 cells showed an ∼ 70% and 100% increase in migration, respectively, upon zeocin treatment when compared to solvent control-treated cells. Interestingly, this phenomenon was not observed with SUM1315MO2 cells. Indeed, zeocin caused a slight decrease in the motility of these cells. Although additional cells must be assayed to reach a firm conclusion, this result raises the possibility that cells with high basal HSATII RNA levels might be less sensitive to zeocin treatment.

**Fig. 8.**
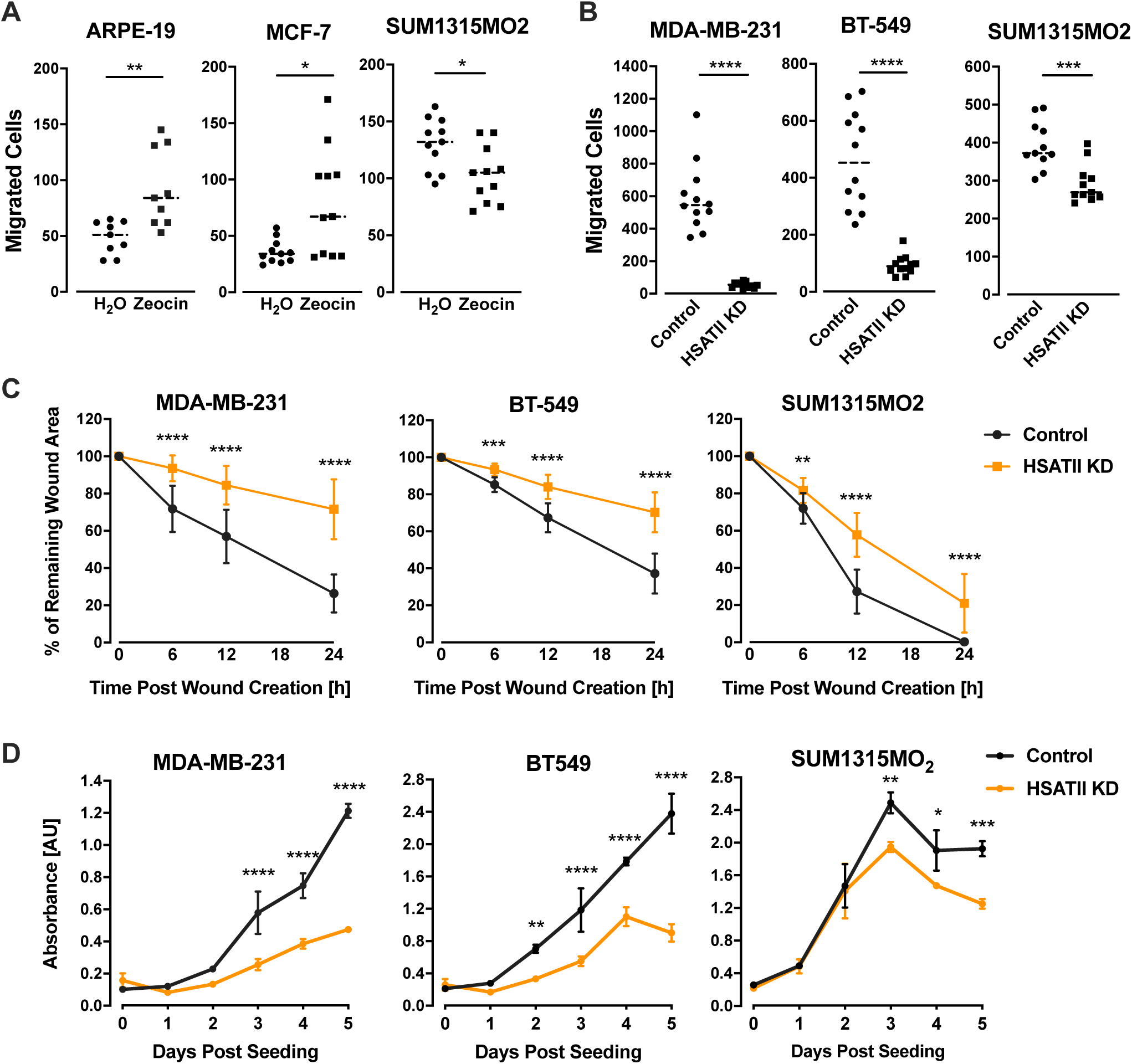
DNA damage-induced HSATII RNA enhances motility and proliferation of breast cancer cells. (*A*) Zeocin treatment increases migration of cells with low basal levels of HSATII RNA. ARPE-1, MCF-7 and SUM315MO2 cells were treated with zeocin (200 μg/mL) for 24 h. Cells were transferred onto transwell inserts, and 24 h later, migrated cells were washed, fixed and nuclei stained. The graph presents a number of cells (with their indicated means) that migrated through a transwell per a field of view (FOV) from biological replicates. *n*=11. (*B*) Transwell assays demonstrated that HSATII KD reduces migration of breast cancer cells. MDA-MB-231, BT-549 and SUM1315MO2 cells were transfected with NT-LNA or HSATII-LNAs. After 48 hpt, equal number of cells was transferred onto transwell inserts. 24 h later, migrated cells were washed, fixed and nuclei stained. The graph presents a number of cells (with their indicated means) migrated through a transwell per FOV from biological replicates. n=12. (*C*) Wound-healing assays demonstrated that HSATII KD reduces migration of breast cancer cells. MDA-MB-231, BT-549 and SUM1315MO2 cells were transfected with NT-LNA or HSATII-LNAs. After 48 hpt, wounds were created and their closure was monitored at indicated times. Data from biological replicates are presented as a percent of remaining wound width mean ± SD. n=20. (*D*) HSATII RNA KD reduces the rate of breast cancer cell proliferation in 2D culture. MDA-MB-231, BT-549 and SUM1315MO2 cells were transfected with NT-LNA or HSATII-LNAs. After 48 hpi, equal number of cells were seeded and cell proliferation was monitored at the indicated times. Cell proliferation is presented as an increase in a measured absorbance (AU – arbitrary units). n=3. \**P*<0.05, \*\**P*<0.01, \*\*\**P*<0.001, \*\*\*\**P*<0.0001.

As MDA-MB-231, BT-549 and SUM1315MO2 cells are highly metastatic (83, 84), we tested effects of HSATII RNA KD on the migratory abilities of these three TNBC cells using transwell migration and wound healing (85) assays. For the transwell migration assay, MDA-MB-231, BT-549 and SUM1315MO2 cells were transfected with either NT-LNA or HSATII-LNAs and motility was assessed 24 h later (Fig. 8*B*). HSATII-deficient MDA-MB-231, BT-549 and SUM1315 cells exhibited significantly reduced migration relative to control-treated cells. MDA-MB-231 and BT-549 cells were the most sensitive to the treatment with HSATII-LNAs, exhibiting 20- and 5-fold decreases in mean migration, respectively. The wound healing assay provided consistent results (Fig. 8*C*). HSATII-deficient MDA-MB-231, BT-549 and SUM1315 cells were significantly slower in closing wounds compared to control treated cells.

As the aggressiveness of TNBC cells is also related to their high proliferation rates, we tested the effects of HSATII KD on the proliferation of MDA-MB-231, BT-549 and SUM1315MO2 cells (Fig. 8*D*). MDA-MB-231, BT-549 and SUM1315MO2 cells were transfected with either NT-LNA or HSATII-LNAs and 24 h later equal numbers of cells were seeded in 2-dimensional cultures and cell proliferation monitored for 5 consecutive days. Although the cell lines proliferated at different rates, the growth rate was significantly reduced for each cell line following HSATII KD.

Our data suggest that HSATII RNA is an important regulator linking a DDR with processes involved in cancer cell migration and proliferation. Further, at least some DNA-damaging cancer therapeutics, through the induction of HSATII RNA, might enhance the migration and proliferation of tumor cells.

## DISCUSSION

Given the ability of HCMV IE1 and 2 proteins to induce a DDR (42, 43) and HSATII RNA (41), and knowing that DNA damage-inducing treatments stimulate the expression of SINEs (54, 55), we tested the possibility that a chemically-induced DDR can also induce HSATII RNA. Zeocin and etoposide robustly induced HSATII RNA in diploid ARPE-19 epithelial cells (Figs. 1*B*) with kinetics (Figs. 1*C*) similar to those described previously for the induction of the transcript by HCMV infection (41). In contrast, UV-C did not induce HSATII (Fig. 1*A*). UV-C causes helix-distorting lesions in DNA, such as pyrimidine(6-4)pyrimidone photoproducts and cyclobutane pyrimidine dimers (50), that can subsequently be converted into DSBs during DNA replication; whereas zeocin (86), etoposide (87) and HCMV infection (88) directly induce DSBs. Our UV-C assay employed confluent monolayers of cells, conditions that did not favor cellular DNA replication and induction of DSBs, so it is possible that extensive DNA DSBs were not produced during the course of the experiment.

Zeocin induced expression from a similar set of HSATII loci as did HCMV-infection (Fig. 2*B* and *C*), and a subset of genes induced by zeocin and virus infection overlapped and included key elements of DDR signaling cascades. Infection and zeocin treatment induced γ-H2AX foci in diploid epithelial cells, a marker for DSBs (Fig. 4). The ATM kinase is the primary mediator of the response to DSBs (89), and KD of ATM in diploid epithelial cells (Fig. 5*A* and *B*) blocked HSATII RNA induction by the DNA-damaging drugs and infection. Zeocin, etoposide and virus infection each induce a DDR and HSATII RNA, and these responses depend on ATM. However, although ATM protein is clearly required, two different active ATM kinase inhibitors (Ku-55933 and AZ31) failed to block HSATII RNA induction by zeocin (Fig. 5*B*). Moreover, ectopic expression of wild-type ATM or kinase dead ATM restored the ability of ATM-deficient ataxia-telangiectasia A-T(-) cells to efficiently induce HSATII RNA in response to zeocin treatment (Fig. 5*D*). Further, KD of Chk2, which is activated by ATM-mediated phosphorylation, and KD of p53, which is phosphorylated and activated by both ATM and Chk2, had no effect on HSATII RNA induction by zeocin (Fig. 6*A* and *B*). Thus, our data indicate that ATM kinase activity and the canonical ATM-Chk2-p53 signaling cascade are not required for HSATII RNA accumulation. Different signaling events fit with the observation that the ATM kinase-dependent induction of CDKN1A and DINO RNAs occurs much more rapidly than the ATM kinase-independent induction of HSATII in response to zeocin (Fig. 1*D*) or etoposide (Fig. 1*E*).

Even though its kinase activity is not essential, ATM protein (or mRNA) is clearly required for HSATII RNA induction. There is precedent for ATM kinase-independent function. ATM controls mitophagy through its interaction with the Parkin E3 ubiquitin ligase; ATM acts via the protein-protein interaction, independently of its kinase activity (90). We anticipate that the kinase-independent role of ATM in the induction of HSATII RNA might be driven by a similar mechanism.

Overexpression of HSATII RNA has been reported for lung, ovarian, prostate and osteosarcoma tumors as well as tumor-derived cell lines (10, 12, 13). Given our current data revealing the importance of the DDR in HSATII RNA expression and with the known impact of deregulated DDRs and cell cycle checkpoints on oncogenesis (91, 92), we probed the physiological consequences of HSATII RNA expression in cultured tumor cells. We tested several breast cancer cell lines growing in adherent, 2-dimensional culture, and higher endogenous HSATII RNA levels correlated with a highly metastatic phenotype (Fig. 7*A*), and HSATII levels were substantially increased in most cell lines by HCMV infection or zeocin treatment (Fig. 7*B* and *C*). Cells characterized by low basal HSATII RNA expression,i.e. ARPE-19 and MCF-7 cells, showed an enhanced migratory phenotype when exposed to zeocin, while SUM1315MO2 cells with higher uninduced levels of HSATII RNA did not (Fig. 8*A*). Perhaps the non-responsive cells had a sufficiently high level of the satellite RNA before induction. HSATII RNA KD markedly reduced migration (Fig. 8*B* and *C*) and proliferation (Fig. 8*D*) in the three TNBC cell lines tested, suggesting a direct relationship between high levels of the RNA and cell phenotype. These observations argue that higher HSAII RNA levels correlate with enhanced proliferation and migration, characteristics of aggressive, metastatic tumors; and they are consistent with an earlier report showing that HSATII copy number gains are a negative prognostic feature of colorectal cancers (17).

DNA-damaging cancer therapeutics can lead to drug resistance (53, 78) and raise concerns about the development of *de novo* primary tumors (93). The induction of HSATII RNA accumulation by DNA-damaging drugs in cultured cells (Fig 1 and 7) and the correlation between elevated HSATII RNA and enhanced tumor cell proliferation and migration (Fig. 8) prompts us to speculate that DNA-damaging chemotherapeutics have potential to enhance malignancy of tumor cells that survive treatment. This effect could be of long or short duration, depending on how long HSATII RNA remains elevated after drug treatment. Negative consequences could derive from immediate effects of HSATII RNA overexpression, as we have measured here, and perhaps overexpression of other non-coding RNAs. In addition, longer-term effects might result from reverse transcription of the overexpressed RNAs followed by integration and copy number expansion in tumors, as described for HSATII (17) and another repeated element, LINE-1 (94).

HCMV RNAs and proteins are found in a variety of human malignancies, and HCMV-infected cells have many properties suggesting the virus might influence tumor progression (95). The ability of HCMV infection, like radiomimetic drugs, to sponsor a DDR (43, 96) with induction of HSATII RNA in healthy cells (41) (Figs. 1 and 2) and breast cancer cells (Fig. 7*B*) leads to a second speculation. Perhaps HSATII RNA induction contributes to the apparent oncomodulatory activity of the virus. Indeed, numerous viruses, many of which are considered tumor viruses, are known to induce a DDR (42); and, therefore, might also induce HSATII RNA contributing to malignancy. For HCMV, even a limited or abortive viral replication documented in cancer cells (56, 80, 81) might lead, *via* IE1/IE2 protein-mediated HSAII RNA induction, to enhanced migratory and proliferative characteristics of cancer cells.

In sum, this study demonstrates that DNA DSBs induce HSATII RNA via a DDR that requires ATM protein but not its kinase activity, demonstrates a link between expression of the RNA and cellular growth and migration phenotype, and establishes a new paradigm to study the biological consequences of HSATII RNA expression - treatment of normal diploid and tumor cells with DNA-damaging agents.

## METHODS

### Cells, viruses and drugs

Human retinal pigment epithelial (ARPE-19) cells, human foreskin fibroblasts (HFFs) (30) and breast epithelial (MCF-10A) cells as well as breast cancer cell lines: MCF-7, MDA-MB-175VII, MDA-MB-361, and MDA-MB-231 were from the American Type Culture Collection (ATCC). SUM1315MO2 breast cancer cells (97) were generously provided by Stephen Ethier (Medical University of South Carolina). A-T(-) fibroblasts, originally named AT22IJE-T, and A-T(+) fibroblasts with a stably expressed functional ATM were created by the lab of Yosef Shiloh (Sackler School of Medicine) (74, 75) and generously provided by Matthew Weitzman (Perelman School of Medicine University of Pennsylvania). ARPE-19, MCF-7, MDA-MB-175VII, MDA-MB-361, MDA-MB-231 and SUM1315MO2 cells were cultured in 10% FBS/DMEM with added Ham’s F-12 nutrient mixture (Sigma-Aldrich) and supplemented with 1x MEM Non-Essential Amino Acids Solution (ThermoFisher Scientific), 1 mM sodium pyruvate, 1x GlutaMAX and 10 mM Hepes, pH 7.4. HFFs, A-T(-) and A-T(+) cells were cultured in 10% FBS/DMEM. Media were supplemented with penicillin G sodium salt (100 units/ml) and streptomycin sulfate (95 units/ml). IE1 and IE2-expressing ARPE-19 cells were described previously (41).

An infectious BAC clone of HCMV strain TB40/E-GFP (TB40/Ewt-GFP) (98) was electroporated into ARPE-19 cells or HFFs to generate viral progeny that were passaged once more in ARPE-19 cells or HFFs, respectively. TB40/E-GFP-epi designates TB40/E-GFP virus grown in ARPE-19 cells (99). All viral stocks were partially purified by centrifugation through a 20% D-sorbitol cushion in buffer containing 50 mM Tris HCl, 1 mM MgCl_2_, pH 7.2, resuspended in DMEM and stored in aliquots at - 80°C. Infections were performed by treating cells with viral inoculum for 2 h, followed by removal of the inoculum and washing with phosphate-buffered saline (PBS; Sigma-Aldrich) before applying fresh medium. To quantify extracellular virions, media from infected cells was collected and spun for 10 min. at 2000 x g to remove debris. To quantify intracellular virions, infected cells were washed with PBS, scrapped and intracellular virus released by 3 rounds of freezing and thawing and then spun for 10 min. at 2000 x g to remove debris. Viral stocks and sample supernatant were titered by plaque assay.Zeocin (InvivoGen) was dissolved in water and stored at −20°C. Etoposide (Millipore Sigma), Ku-55933 (Tocris) and AZ31 (Selleckchem) were dissolved in DMSO and stored at −20°C.

### RNA sequence analysis (RNA-seq)

RNA was collected in QIAzol Lysis Reagent (Qiagen) and isolated using the miRNeasy Mini Kit (Qiagen). DNA was removed from samples using Turbo DNase (Thermo Fisher Scientific) and RNA quality was assessed using the Bioanalyzer 2100 (Agilent Technologies). Sequencing libraries were prepared using the TruSeq Stranded Total RNA with Ribo-Zero kit (Illumina), and sequenced on Illumina HiSeq2500 sequencer instrument in paired-end, rapid mode (2x 150bp).

For RNA-seq analysis, the Galaxy instance of Princeton University was used (100). RNA-Seq data were de-multiplexed based on indexes. Phred quality scores were checked by using the FastQC toolkit and were greater than 26 for more than 90% of the read length. Human and HCMV fasta and annotation (.gtf) files were created for mapping by combining sequences and annotations from Ensembl annotation, build 38 (GRCh38), Repbase elements (release 19) and TB40 (EF999921) when appropriate. Quality filtered reads were mapped to the concatenated human-virus genomes using RNA STAR (57) (Galaxy version 2.6.0b-2). The featureCounts program (101) (Galaxy version 1.6.4) was used to measure gene expression. The resulting files were used as input to determine differential expression for each gene utilizing DESeq2 (102) (Galaxy version 2.11.40.6). Fold changes in gene expression were considered significant when the adjusted *P* value (q value) for multiple testing with the Benjamini–Hochberg procedure, which controls for the False Discovery Rate (FDR), was *<*0.05. To analyze HSATII expression, aligned reads were assigned using the featureCounts function of Rsubread package (103) with the external HSATII annotation obtained from a RepeatMasker (104) using GRCh38 and Repbase consensus sequences. This produced the raw read counts. HSATII expression in terms of CPM (counts per million reads) was computed and normalized across samples using the trimmed mean of M-values method (TMM) (105). To calculate the percent of HSATII reads originating from each chromosome, we identified uniquely mapped reads that exclusively overlapped with HSATII repeat. The number of normalized counts of HSATII reads mapped to each chromosome was computed and the percentage of these reads mapping to each chromosome was calculated as described previously (41). To visualize read coverage tracks for alignment files, RNA STAR-generated BAM files were imported into the Integrative Genomics Viewer (106) (IGV; version 2.6.3) and aligned with human genome assembly GRCh38 focused on the HSATII locus on chromosome 16. The scale of read coverage peaks was adjusted for experimentally paired samples. To create Venn diagrams, based on data from the DESeq2 analysis the list of significantly expressed genes in HCMV-infected or zeocin-treated cells compared to control cells were imported into a web-based tool InteractiVenn (107). Gene Set Enrichment Analysis (GSEA) (58) was performed on the list of common genes differentially expressed in both HCMV-infected and zeocin-treated cells using the hallmark gene set of the Molecular Signature Database (MSigDB) (59). A matrix of differentially expressed genes from the data set significantly matching the hallmark gene set of MSigDB was composed and a list of the hallmark gene subsets was ordered based on a number of overlapping genes, *p* value determining the probability of association with a given gene set and a FDR q-value.

### Quantification of nucleic acids

For analysis by qRT-PCR, RNA was extracted from samples collected in QIAzol lysis reagent using the miRNeasy kit (Qiagen). cDNA was made from 1 µg of total RNA with random hexamer primers and MultiScribe reverse transcriptase (Applied Biosystems), according to the manufacturer’s protocol. Transcript-specific primer sequences used in the study were either previously reported (41) or newly designed: CDKN1A Forward 5’-TTTAACAGCCTGCTCCCTTG-3’, Reverse 5’-AGTTTGCAACCATGCACTTG-3’; DINO Forward 5’-GGAGGCAAAAGTCCTGTGTT-3’, Reverse 5’-GGGCTCAGAGAAGTCTGGTG-3’ (49); E2F1 Forward 5’-CCGTGGACTCTTCGGAGAAC-3’, Reverse 5’-ATCCCACCTACGGTCTCCTC-3’; E2F2 Forward 5’-TGGGTAGGCAGGGGAATGTT-3’, Reverse 5’-GCCTTGTCCTCAGTCAGGTG-3’; E2F3a Forward 5’-TTTAAACCATCTGAGAGGTACTGATGA-3’, Reverse 5’-CGGCCCTCCGGCAA-3’ (108); E2F3b Forward 5’-TTTAAACCATCTGAGAGGTACTGATGA-3’, Reverse 5’-CCCTTACAGCAGCAGGCAA-3’ (108); E2F4 Forward 5’-TGGAAGGTATCGGGCTAAT-3’, Reverse 5’-CAATCAGTTTGTCAGCAATCTC-3’; E2F5 Forward 5’-CGGCAGATGACTACAACTTTA-3’, Reverse 5’-GATAACAGTCCCAAGTTTCCA-3’; ATM Forward 5’-GCACTGAAAGAGGATCGTAAA-3’, Reverse 5’-GAGGGAACAAAGTCGGAATAC-3’; ATR Forward 5’-ATATCACCCAAAAGGCGTCGT-3’, Reverse 5’-TGCTCTTTTGGTTCATGTCCAC-3’; PRKDC Forward 5’-AAATGGGCCAGAAGATCGCA-3’, Reverse 5’-AGGTCCAGGGCTGGAATTTT-3’; CHEK1 Forward 5’-TTTGGACTTCTCTCCAGTAAAC-3’, Reverse

5’-GCTGGTATCCCATAAGGAAAG-3’; CHEK2 Forward 5’-CGCGGTCGTGATGTCTCGGG-3’, Reverse 5’-CGCTGCCATGGGGCTGTGAA-3’; TP53 Forward 5’-AGGGATGTTTGGGAGATGTAAG-3’, Reverse 5’-CCTGGTTAGTACGGTGAAGTG-3’ (109). qPCR was performed using SYBR Green master mix (Applied Biosystems) on a QuantStudio™ 6 Flex Real-Time PCR System (Applied Biosystems). Transcript levels were analyzed using the delta delta Ct method and GAPDH was used as an internal control (110). Error ranges are reported as standard deviation of the mean (SD).

### Analysis of proteins

For Western blot analysis, transfected A-T(-) were harvested using lysis buffer (50 mM Tris-HCl, pH 7.5, 5 mM EDTA, 100 mM NaCl, 1% Triton X-100, 0.1% SDS, and 10% glycerol). Samples were mixed with 6xSDS sample buffer (325 mM Tris pH 6.8, 6% SDS, 48% glycerol, 0.03% bromophenol blue containing 9% 2-mercaptoethanol). Proteins were separated by electrophoresis (SDS-PAGE) and transferred to ImmunoBlot polyvinylidene difluoride (PVDF) membranes (BioRad Laboratories). Western blot analyses were performed using primary antibodies recognizing ATM (D2E2; Cell Signaling Technology), GFP (D5.1; Cell Signaling Technology), p-Chk2 (T68) (2661; Cell Signaling), Chk2 (2662; Cell Signaling) and β-actin-HRP (Abcam). Donkey anti-rabbit (GE Healthcare Biosciences) conjugated with horseradish peroxidase was used as a secondary antibody. Western blots were developed using WesternSure ECL Detection Reagents (Licor).

### Immunofluorescence

For immunofluorescence assays, ARPE-19 cells grown in 96-well plates were infected with TB40/E-GFP-epi (3 IU/cell), treated with zeocin (200 µg/mL) or left untreated. After 24 h, cells were fixed in 100% methanol (Sigma) and incubated in blocking solution [PBS with 3% bovine serum albumin prior to the addition of a primary mouse monoclonal antibody specific for phospho-Histone H2AX (Ser13; clone JBW301; Millipore Sigma). Following incubation, cells were washed with PBS, and a donkey Alexa488-conjugated anti-mouse secondary antibody (Santa Cruz Biotechnology) was applied. Nuclei were counterstained with the Hoechst stain. Cells were visualized and % of γ-H2AX-positive cells was calculated using the Operetta high-content imaging and analysis system (Perkin Elmer) with 20X objective.

### Plasmid transfection

Plasmids used in the study: pCMV-Neo-Bam was a gift from Bert Vogelstein (Addgene plasmid #16440). pCMV-Neo-Bam E2F3a and pCMV-Neo-Bam E2F3b were gifts from Jacqueline Lees (Addgene plasmid #37970 and #37970, respectively). pCMVHA E2F1 and pCMVHA E2F2 were gifts from Kristian Helin (Addgene plasmids #24225 and #24226, respectively). ARPE-19 cells at 70-80% confluency were transfected with 1 µg of each plasmid using Lipofectamine 3000 (Thermo Fischer Scientific) according to the manufacturer’s instructions. 72 h later, plasmid-transfected cells were harvested using TRIzol and samples were stored at −80°C.

### Cell stress induction

For the induction of stress by serum deprivation (111), cells were washed with 1xPBS, fed with medium lacking FBS and cultured for 24 h. For the oxidative stress (112), cells were washed with 1xPBS, fed with medium lacking FBS, but containing 100 μM of H_2_O_2_ and cultured for 24 h. For UV-irradiation-based stress (113), cells were washed with 1xPBS, exposed to the ultra violet light (50 J/m^2^) using a Stratalinker (Stratagene), washed with 1xPBS, supplemented with 10%FBS/DMEM and cultured for 24 h.

### Locked nucleic acid and siRNA KD

Locked nucleic acid (LNA™; Exiqon, Skelstedt, Denmark) oligonucleotides targeting HSATII RNA were described previously (41). Control NT-LNA or HSATII-LNAs (100-200 nM) were transfected into cells using Lipofectamine RNAiMAX® Reagent (Thermo Fisher Scientific) according to the manufacturer’s instructions. LNA-treated ARPE-19 cells were incubated for 24-48 h before being subjected to subsequent experimental procedures.

siRNAs targeting E2F1 (E2F1 siRNA), E2F2 (E2F2 siRNA), E2F3 (E2F3 siRNA), E2F4 (E2F4 siRNA), E2F5 (E2F5 siRNA), CHEK1 (CHEK1 siRNA) and CHEK2 (CHEK2 siRNA) were siGENOME SMARTpools (Dharmacon). siRNAs targeting ATM (ATM siRNA), ATR (ATR siRNA), PRKDC (PRKDC siRNA) and TP53 (TP53 siRNA) were ON-TARGETplus SMARTpools (Dharmacon). All siRNAs and appropriate NT siRNAs (siGENOME Non-targeting siRNA Pool #1 and ON-TARGETplus Non-targeting Control Pool; Dharmacon) were purchased from Life Technologies. ARPE-19 cells were grown to ∼80% confluence and then transfected with siRNA using Lipofectamine® RNAiMAX Reagent (Life Technologies). Cells transfected with non-specific, scrambled siRNA (NT siRNA) (Life Technologies) served as a negative control. Following 48 h incubation with siRNA, cells were washed with 1xPBS and subjected to subsequent experimental procedures.

### Cell migration, proliferation and toxicity assays

Wound healing and transwell migration assays were performed as described previously (41). For wound healing assays, confluent monolayers of NT- LNA- or HSATII-LNA-transfected MDA-MB-231, BT-549 and SUM1315MO2 cells were scratched with a 1-ml pipet to create wounds. The process of wound closure was monitored in time using a Nikon Eclipse TE2000-U inverted microscope. The average wound width (in arbitrary units) of ARPE-19 cells was calculated from 6 measurements for each experimental arm from the captured images using ImageJ software (114). Results are plotted as a mean percent of remaining wound width (SD). For transwell assays, NT-LNA- or HSATII-LNA -transfected MDA-MB-231, BT-549 and SUM1315MO2 or zeocin-treated ARPE-19, MCF-7 and SUM1315MO2 cells were trypsinized and 50,000 cells were seeded onto each filter in FBS-free medium containing ITS Liquid Media Supplement (Sigma-Aldrich). After 24 h, filters were washed, cells were fixed and migrated cells on the bottom surface of the filter were stained with 0.2% crytal violet solution. Migrated cells were imaged and quantified using ImageJ software (114).

To monitor proliferation, cells were transfected with NT-LNA or HSATII-LNAs. After 48 hpt, equal number of cells (2500 cells/well) were seeded into 96-well plates and cultured in 10% FBS/DMEM with added Ham’s F-12 nutrient mixture and supplemented with 1x MEM Non-Essential Amino Acids Solution, 1 mM sodium pyruvate, 1x GlutaMAX and 10 mM Hepes. Cell numbers were quantified using a colorimetric 3-(4,5-dimethylthiazol-2-yl)-5-(3-carboxymethoxyphenyl)-2-(4-sulfophenyl)-2H-tetrazolium (MTS)-based CellTiter 96® AQ_ueous_ One Solution Cell Proliferation Assay (Promega, Madison, WI) according to the manufacturer’s protocol. Absorbance was measured at 490 nm using the SpectraMax Plus 384 Microplate reader (Molecular Devices).

To assay toxicity, ARPE-19 cells were treated with zeocin, etoposide, Ku-55933 or AZ31 at specified concentrations for 24 h or with water or DMSO as solvent controls. Each tested concentration of a specific drug contained the same concentration of solvent. At indicated times, the CellTiter 96® AQ_ueous_ One Solution Cell Proliferation Assay (Promega) was performed according to the manufacturer’s instructions. Absorbance was measured at 490 nm using the SpectraMax Plus 384 Microplate reader (Molecular Devices).

### Statistical analysis

To determine statistical significance between two conditions in experiments, unpaired, two-tailed *t*-tests with Welch’s correction were performed, otherwise one-way ANOVA was performed between the arrays of data from distinct samples to determine *P* values. *P* value <0.05 was considered significant. Significance is shown by the presence of asterisks above data points with one, two, three or four asterisks representing *P*<0.05, *P*<0.01, *P*<0.001 or P<0.0001, respectively. Only significant *P* values are reported.

### Data availability

Raw RNA-seq data will be available from the NCBI Gene Expression Omnibus (GEO). All relevant experimental data are available from the authors.

## ACKNOWLEDGMENTS

We thank Stephen Ethier (Medical University of South Carolina) for generously providing SUM1315MO2 cells; Matthew Weitzman (Perelman School of Medicine, University of Pennsylvania) for generously providing A-T(-) and A-T(+) fibroblasts, Adam Oberstein (University of Illinois) for creating IE1/IE2-expressing ARPE-19 cells, Alexander Solovyov (Memorial Sloan Kettering Cancer Center) for advice and assistance with RNA-seq analyses and members of the T.S. laboratory for scientific discussions.

This work was supported by National Institutes of Health Grants AI112951 and AI142520. M.T.N. was partially supported by American Cancer Society Fellowship PF-14-116-01-MPC.

## AUTHOR CONTRIBUTIONS

M.T.N. designed and performed experiments, analyzed the data and wrote the manuscript. T.S. supervised the study, analyzed the data and edited the manuscript.

